# The micro-plant remains preserved on lithics and in the associated sediment of the Hoabinhian Laang Spean cave site, Battambang province, Cambodia

**DOI:** 10.64898/2026.07.26.740757

**Authors:** Céline E. Kerfant, Ruxi Yang, Ignacio Clemente-Conte, Sophady Heng, Isabel Expósito, Antonio Pérez-Balarezo, Simon Puaud, Valery Zeitoun, Cyril Viallet, Justin Guibert, Hubert Forestier

## Abstract

This study seeks to advance understanding of how ancient Hoabinhian communities engaged with the diversity of their environments, focusing on plant resource management and the role of plants in technological and subsistence strategies at the Laang Spean site, Cambodia. The methodological approach relies on phytolith assemblages recovered from sediments associated with micro-plant residues adhering to the lithic tools, interpreted using anatomical and histological knowledge of plant tissues.

Phytolith extraction from nine stratigraphic layers revealed distinct differences between Hoabinhian contexts (Stratigraphical Unit=SU): the layer 9 contained mainly palm trees phytoliths, while layers 10 and 11 showed higher phytolith diversity and diagnostic grass forms.

Besides the Monocots strong assemblage results between the two main periods studied, a diversity of phytolith arboreal taxa indicate broader plant use and environmental complexity during Hoabinhian occupations (consistent with the other archaeobotanical studies held on Hoabinhian archaeological sites in the region).

Upon the nine lithics artefacts studied, only one has delivered micro-plant remains identified as leaf-culm part of Monocots (plants that do not produce wood), and most probably of bamboo origin. This multidisciplinary research aims to refine interpretations of Hoabinhian plant exploitation (subsistence and environmental context), and contributes to current debates on “forest technologies” in Southeast Asia.

## Introduction

The Hoabinhian is a major lithic technocomplex in the later prehistory of (sub)tropical Asia, documented from at least ca. 43.5 ka at Xiaodong Rockshelter in Yunnan (Ji et al., 2016) through the Middle Holocene (Forestier et al. 2013), across a vast region from southwestern China to Sumatra (Forestier et al., 2013; Shoocongdej, 2022; Forestier, 2024; Wu et al., 2024).

It has been variously conceptualised as an ethnic label, a chronological unit, a subsistence strategy and a lithic technocomplex, reflecting long-standing debates over its definition and internal variability (Gorman, 1969, 1971; Glover, 1977; Van Tan, 1994; Moser, 2001; Forestier et al., 2017, 2021, 2025; Chitkament, 2023; Marwick, 2017; Zeitoun et al., 2019a; Zhou et al., 2025).

Developed across highly diverse and fluctuating ecosystems over a long time span, this technocomplex raises questions about the variability of Hoabinhian groups and the adaptive strategies they used to manage different environments (Maxwell, 1999; Zeitoun et al., 2024; Forestier et al., 2025).

The use of plants is key to understanding the technological and cultural complexity of Hoabinhian societies, yet this aspect remains poorly documented due to the inherent degradability of plant materials, particularly in tropical environments (Bannanurag, 1988; Marwick, 2007; Zeitoun et al., 2008; Sémah & Sémah, 2012; Forestier et al. 2015, 2017, 2021; Heng et al., 2016). Although archaeobotanical research on Hoabinhian sites has been limited, some key results have emerged, attesting to the rich potential of such studies for the region (Table 1). Carpological and other charred-remain analyses highlight the importance of plant resources in Hoabinhian assemblages, for example at Spirit Cave in Thailand between 12,000 and 7,000 BP (Gorman, 1971; Bulbeck & Marwick, 2022). In Vietnam, between 18,000 and 7,000 BP, numerous sites have yielded information on plant use through combined pollen and carbonised remains; at Xom Trai, carpological analysis identified ten wild forest taxa, including *Aeolocarpus* sp., *Gnetum* sp. and several palms (Nguyen, 2008). A recent carpological study of Hiem Cave in northern Vietnam (8,500–8,200 BP) highlighted the management of tropical plant taxa such as *Canarium* sp., while analyses of charred remains from Mai Da Dieu, Dong Cang and Con Moong document alternating wet and dry episodes through two distinct signatures of forest tree groups with *Juglans*-like sp. and *Quercus* sp. (Masojć et al., 2023). Taken together, these findings show that a wide diversity of tropical plant species played a significant role in Hoabinhian subsistence, including both hardwood taxa and various monocots (e.g. taro, bamboo and palms).

**Table 1.**
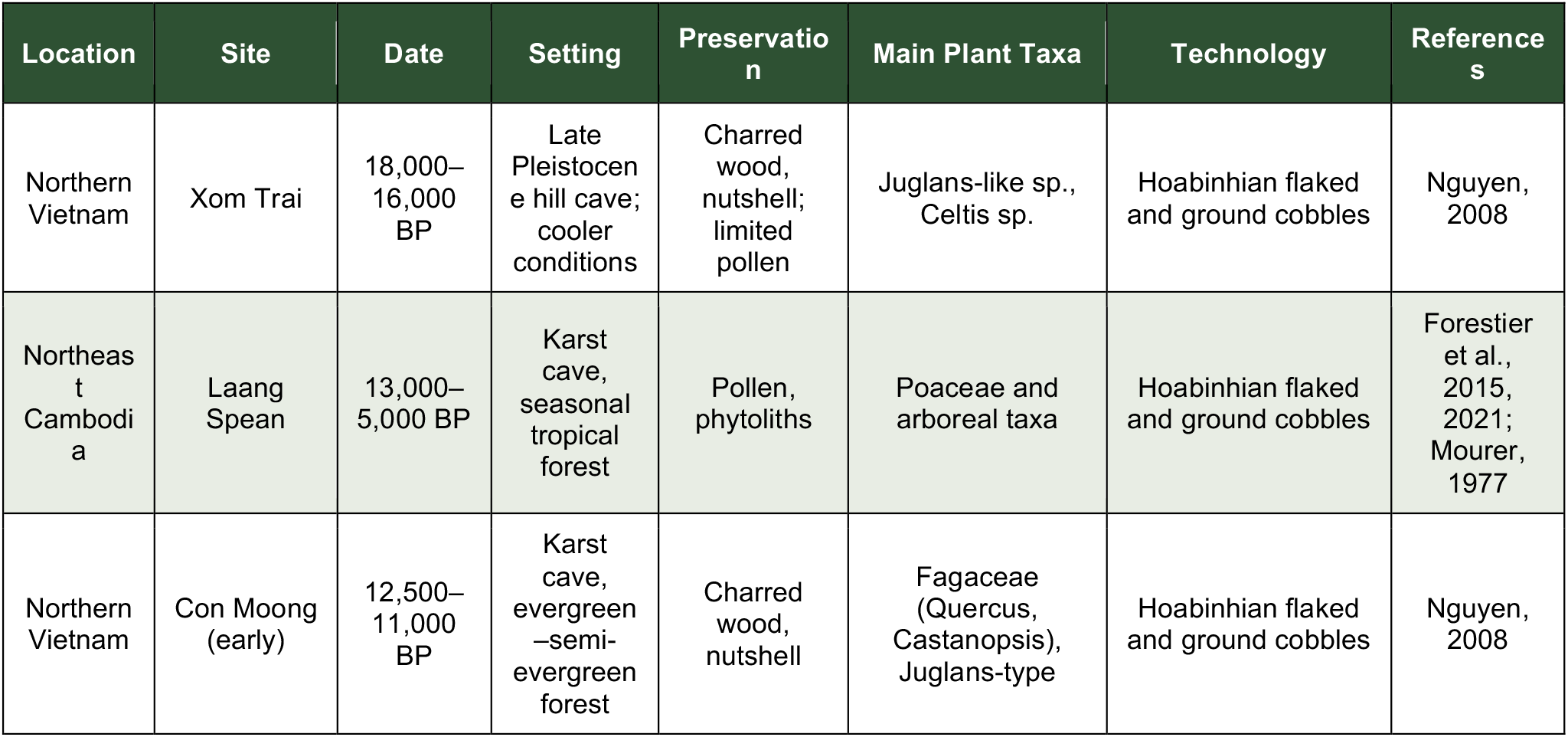
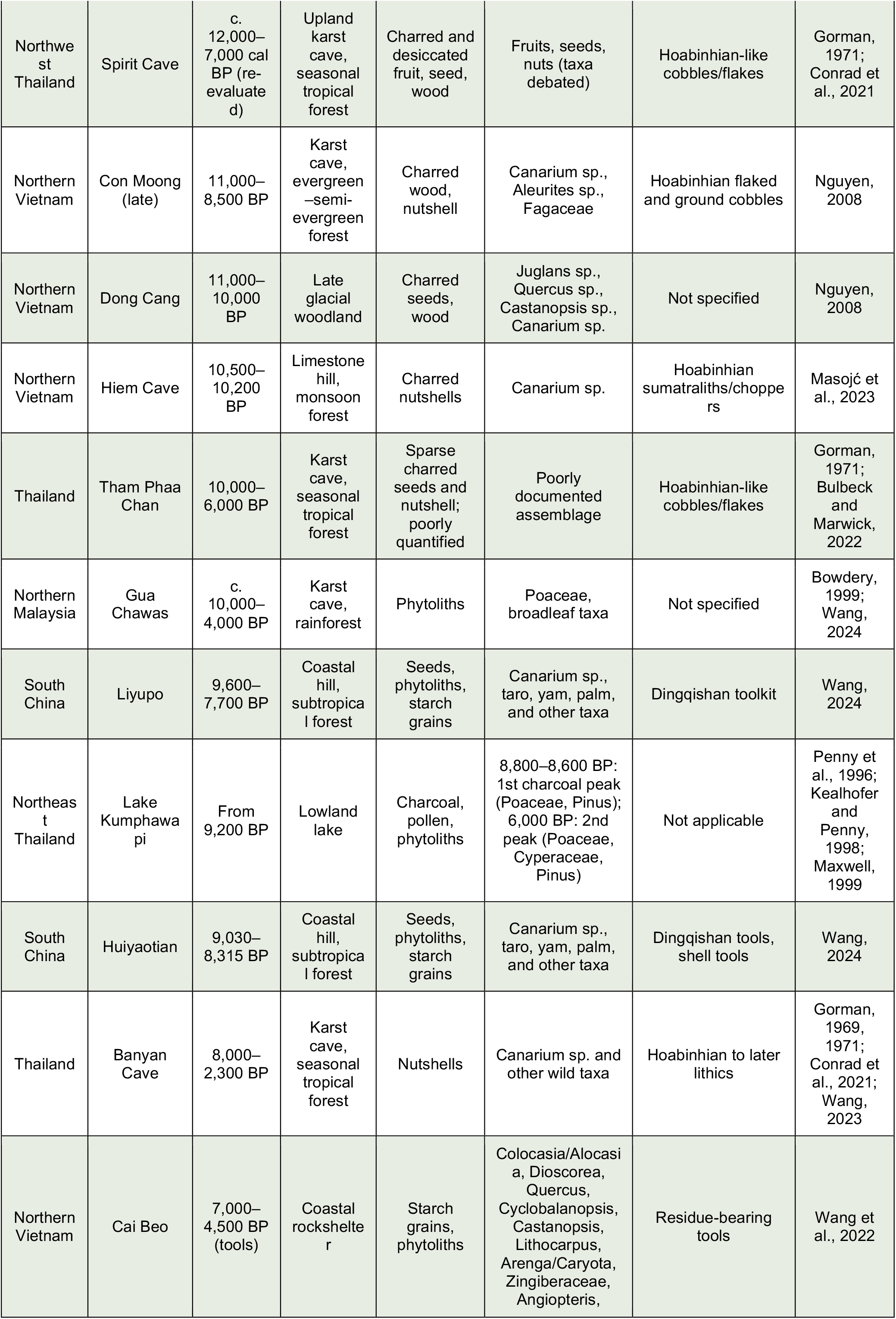
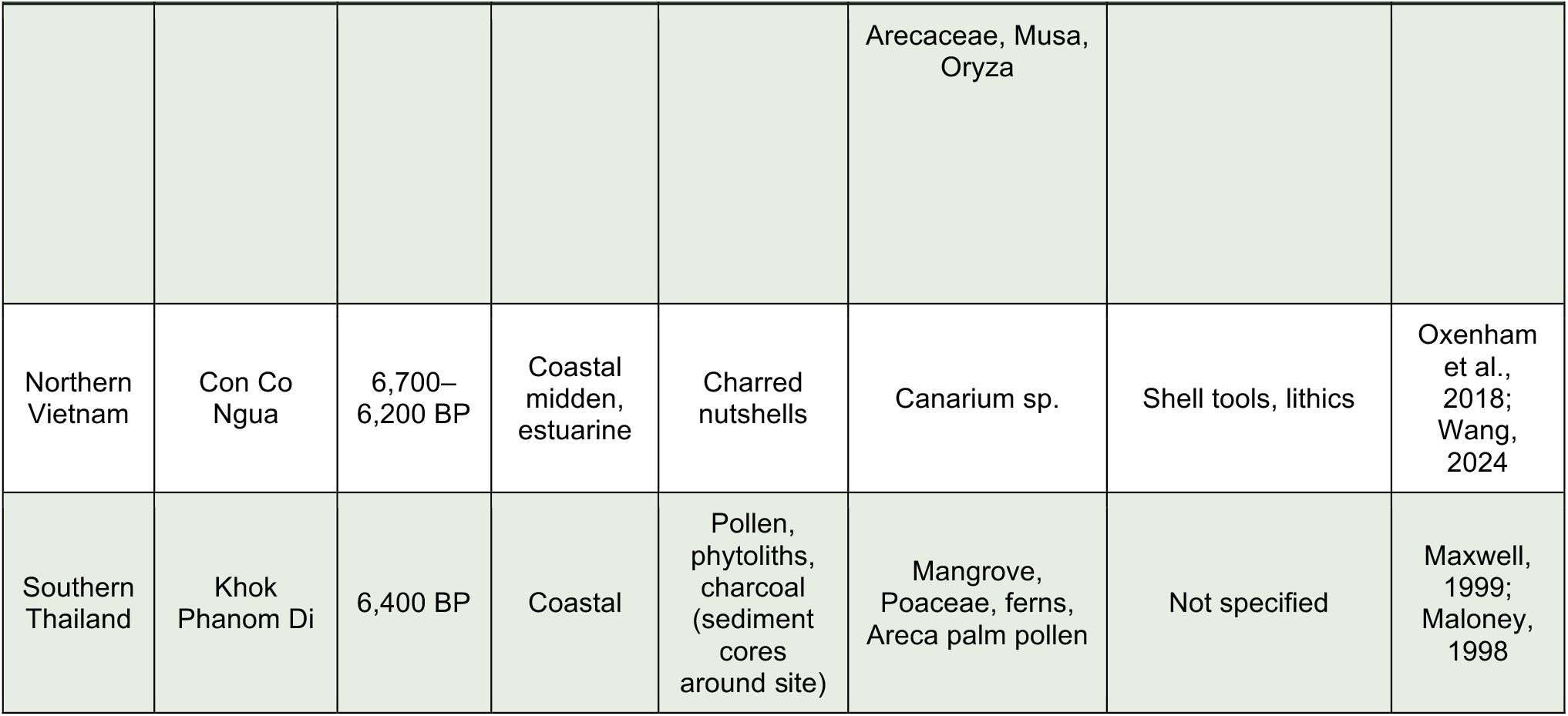
Chronological overview of archaeobotanical evidence associated with lithic assemblages from regional sites spanning the early to late Pleistocene. Sites arranged in chronological order (oldest to youngest) by the start of their reported date range, in years Before Present (BP).

The analysis of phytoliths, silica-based plant microremains, has become a crucial tool for identifying plant use in deep-time archaeological contexts. These can be retrieved from a variety of archaeological contexts and often persist long after other plant tissues have decomposed. Therefore, phytoliths have been used to reveal plant presence and use in Southeast Asia and beyond. They are offering valuable insights into the exploitation of monocots (plants that do not produce wood and which are generally prolific phytolith morphotype producers), such as early banana cultivation at Fahien Rock Shelter in Sri Lanka (Premathilake & Hunt, 2018) and rice in Thailand (Kealhofer & Piperno, 1994). Phytolith evidence for early plant practices, particularly millet and rice cultivation in different ecological and cultivation regimes, is especially informative, as experimental work shows that water stress affects both the quantity and morphology of phytoliths produced by rice (Weisskopf et al., 2015). In cave settings, phytolith analysis is particularly valuable because plants generally cannot grow naturally inside caves, so their microremains are more likely to derive from human activity (Paz, 2005).

Plant microremains can also be resilient when preserved on lithic tools; residues tend to mineralise, allowing recovery of phytoliths and other microbotanical remains directly from tool surfaces (Hardy et al., 2020). This integrative plant-anatomy approach remains under-exploited, yet it can provide substantial information on plant taxonomy and the specific plant parts processed (Litzses-Sabo, 2019). Recent work therefore opens promising avenues for linking stone-tool function with plant exploitation in tropical archaeological contexts (Fuentes et al., 2020; Xhauflair et al., 2023, 2024).

The aim of this article is to contribute to these debates by examining the Hoabinhian technocomplex at Laang Spean Cave through an integrated analysis of lithic technology and plant remains. Specifically, we investigate what combined evidence from lithic and plant materials reveals about the complexity and diversity of material culture within Hoabinhian assemblages. To address this question, we apply plant anatomical and phytolith analyses to identify micro botanical remains preserved on a selected sample of lithic tools (n = 9). We then explore the relationships between tool morpho-structural characteristics, use-wear signatures, and plant-processing activities, including food preparation, raw material working, and the production of hafted or composite tools.

## Material and methods

### Materials

Laang Spean Cave, located in Battambang Province, northwestern Cambodia, is one of the most significant Hoabinhian sites in Mainland Southeast Asia (Fig. 1). Excavated since the French colonial era in the 1960s by the Mourer’s, the site has been re-investigated by the French-Cambodian Prehistoric Mission since 2009, yielding new stratigraphic, chronocultural, anthropological, paleontological and archaeozoological data.

**Figure 1.**
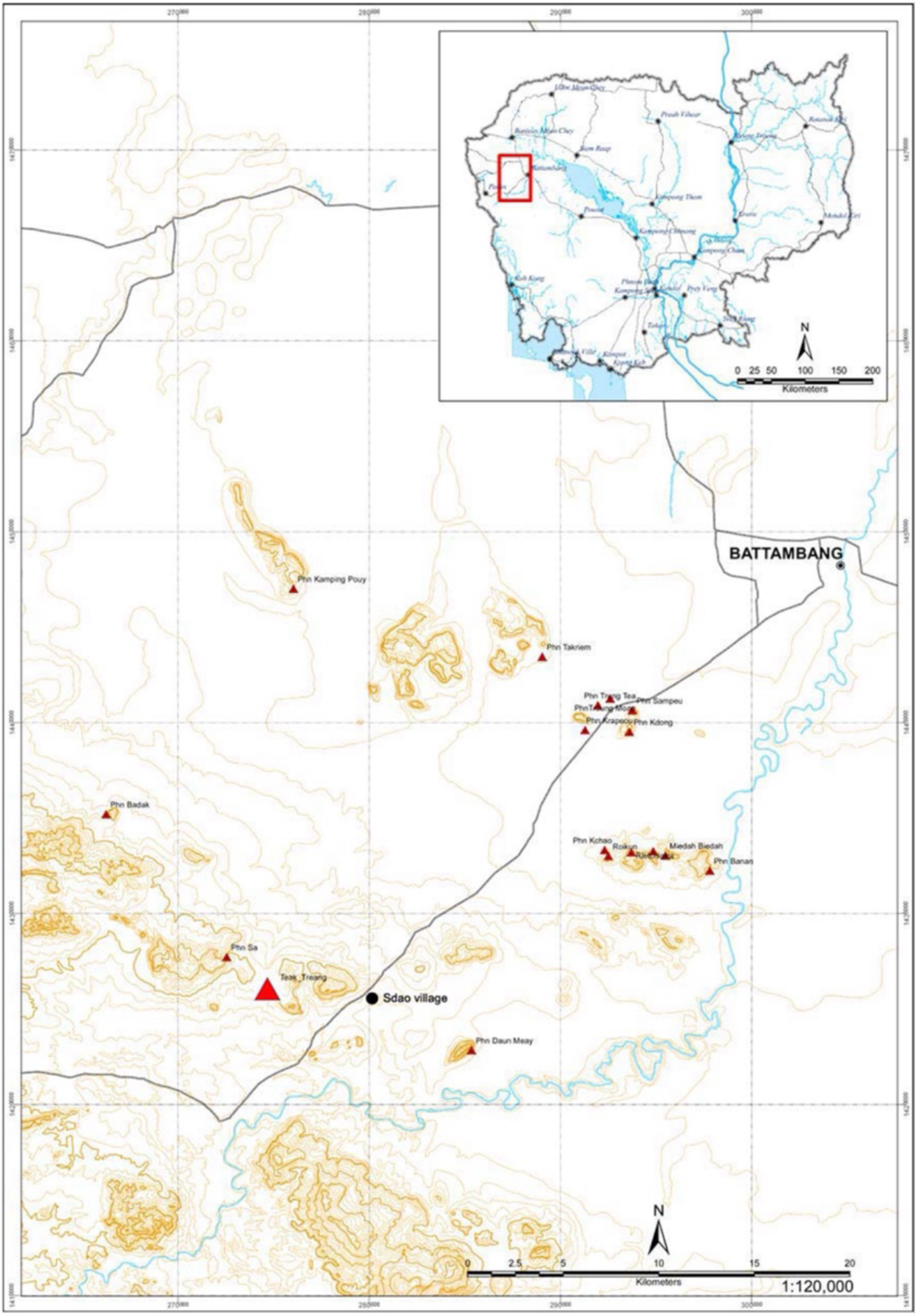
Situation of Laang Spean Prehistoric cave site, Northwest Cambodia in the plain of Battambang (From Heng et al., 2016).

Hoabinhian occupation is dated to approximately 13,000–5,000 cal. BP between SU9 to SU11b1 (Forestier et al. 2021). Beneath Level 11b lies the underlying “pre-Hoabinhian” level, characterised by a flake industry made from chert nodules and dated to the post-LGM period (<21 ka) (Fig.2). A lower archaeological level has been recorded at a depth of 5 m and is recently dated to 100 ka BP (Guibert et al., in review). This long occupational sequence provides a rare opportunity to study technological and subsistence transitions from mobile hunter-gatherer groups to more sedentary communities in the region (Heng et al., 2016; Zeitoun et al., 2024). At Moh Khiew Cave (southern Thailand), the lithic assemblage is dominated by small chert flake tools, including “slug” pieces and occasional bifacial elements, and does not include the cobble tools typically associated with the Hoabinhian tradition (Forestier et al., 2015). Similarly, the Doi Pha Kan site in Lampang Province (northern Thailand) presents a lithic assemblage based on the exploitation of siliceous slabs to produce composite asymmetrical tools, differing from more conventional Hoabinhian lithic forms (Forestier et al., 2025).

For the present study, nine sediment samples were collected from distinct stratigraphic units (Fig. 2) by Hubert Forestier and Sophady Heng, directors of the French-Cambodian Prehistoric Mission, for phytolith analysis. In addition, nine lithic artefacts were selected for microbotanical analysis by one of us (I. Clemente-Conte), a use-wear specialist, based on their morphological variation, technological relevance and potential for preserving plant residues.

**Figure 2.**
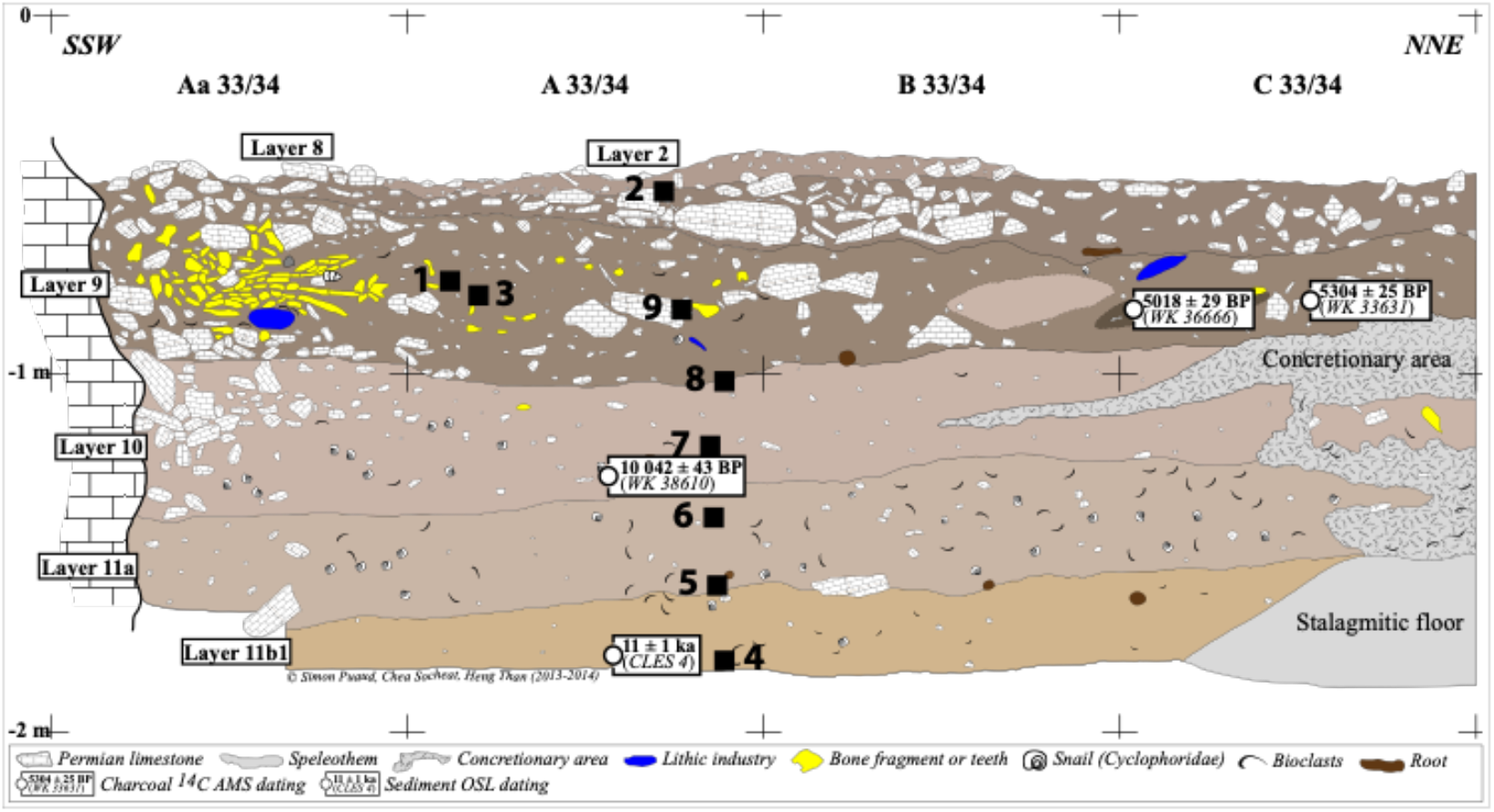
Section showing the Hoabinhian level and dating of Laang Spean, Cambodia (after S. Puaud in: Forestier et al. 2015, 2021). The squares indicate the analysed samples.

## Methodology

The research combines lithic use-wear and residue analyses with sediment phytolith study to explore the relationships between tool function, plant exploitation, and local vegetation (Kealhofer et al, 1999).

For this, phytoliths from the nine Hoabinhian layers were extracted (Fig. 2) following the protocol of Albert & Weiner (2001) at the Archaeobotany Laboratory of IPHES-CERCA, Tarragona. Microscope slides were prepared using Entellan™ New mounting medium. Phytoliths were identified and counted under transmitted white light with an Iscope-Euromex compound light microscope at 400× magnification at the Archaeobotany Unit, IPHES-CERCA. For each slide, three transects were counted, after which the entire slide was examined to locate silica skeletons. Representative phytoliths and silica skeletons were photographed using the software DpxViewPro. We used ICPN 2.0 nomenclature to classify the phytoliths (ICPT, 2019).

Micro-plant remains potentially preserved on the lithic artifacts were examined with a digital Hirox KH-8700 microscope at IPHES, following a non-destructive protocol that combines low-magnification use-wear analysis (around 50×) with high-resolution microwear observation (100–400×) (Hayes et al., 2017). This approach enables the documentation of both macro-and micro-traces with the same instrument, allowing continuous observation from residue mapping to detailed characterization and, when possible, taxonomic assessment of plant remains. Digital microscopy thus serves as an effective bridge between lithic and plant studies, supporting integrative interpretations of tool function and plant use (Hardy et al. 2020; Martín-Viveros & Ollé, 2020; Kerfant, 2022; Xhauflair et al., 2023).

## Results

### Phytoliths from sediment

Sample 2 contained very few phytoliths, and Sample 8 was lost during the heating step of the extraction process; both were excluded from further analysis.

#### SU 9 (4258–3632 cal BP)

Layer 9 (Samples 1, 3, and 9) is characterized by the presence of spheroid echinate (SPH_ECH) phytoliths (Table 2). Spheroid psilate (SPH_PSI) morphotypes, occurring either within globular forms attributed to palm or as isolated morphotypes, were identified only in this layer (Figure 3). These morphotypes are not present in the SU 10–11.

**Table 2.**
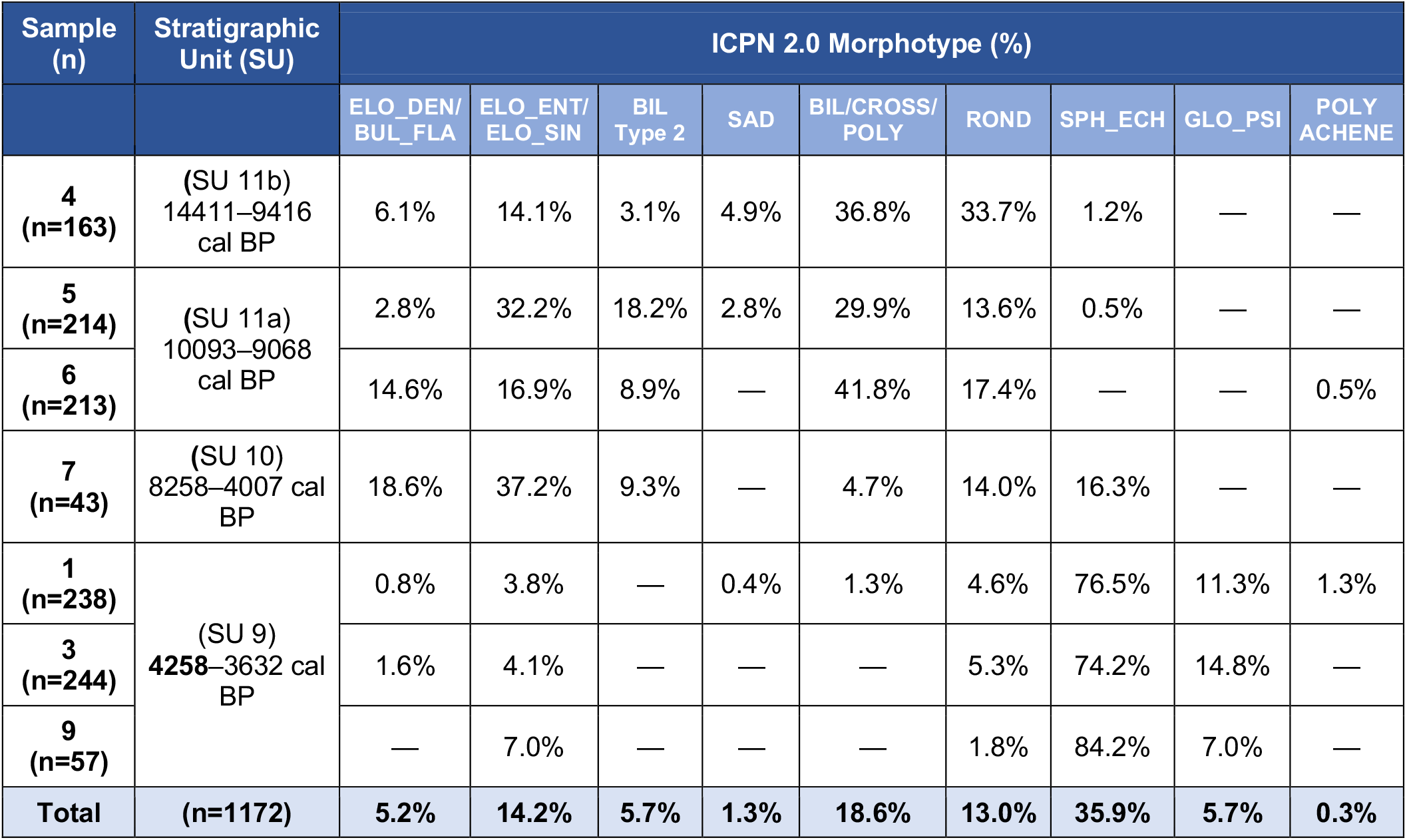
Phytolith morphotype percentages by sample and stratigraphic unit. Samples are ordered chronologically from oldest to youngest according to stratigraphic unit (cal BP), following Antonio Perez Balarezo’s recent calibration work. A dash (—) indicates 0% or not observed.

**Figure 3.**
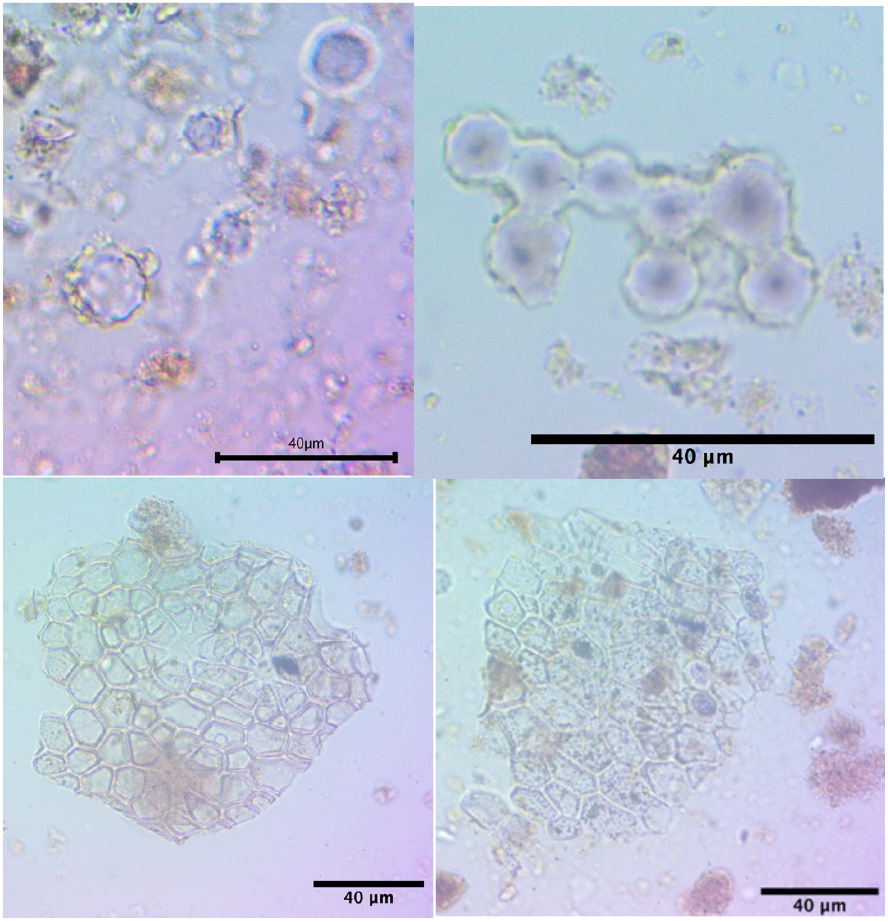
Phytoliths from SU 9 (from top left to bottom right : Spheroid echinate (SPH_ECH), Spheroid psilate (SPH_PSI) and skeletons.

#### SU 10 (8258-4007 cal BP), SU 11a (10,093-9068 cal BP), and SU 11b (14,4119-416 cal BP)

Layers 10 and 11 (Samples 4 to 7) are characterised by higher phytolith density and diversity, with assemblages dominated by grass silica short-cell phytoliths (GSSCP) attributable to Poaceae (Figure 4). In contrast to SU 9, SPH_ECH morphotypes are rare or absent (Table 2).

**Figure 4.**
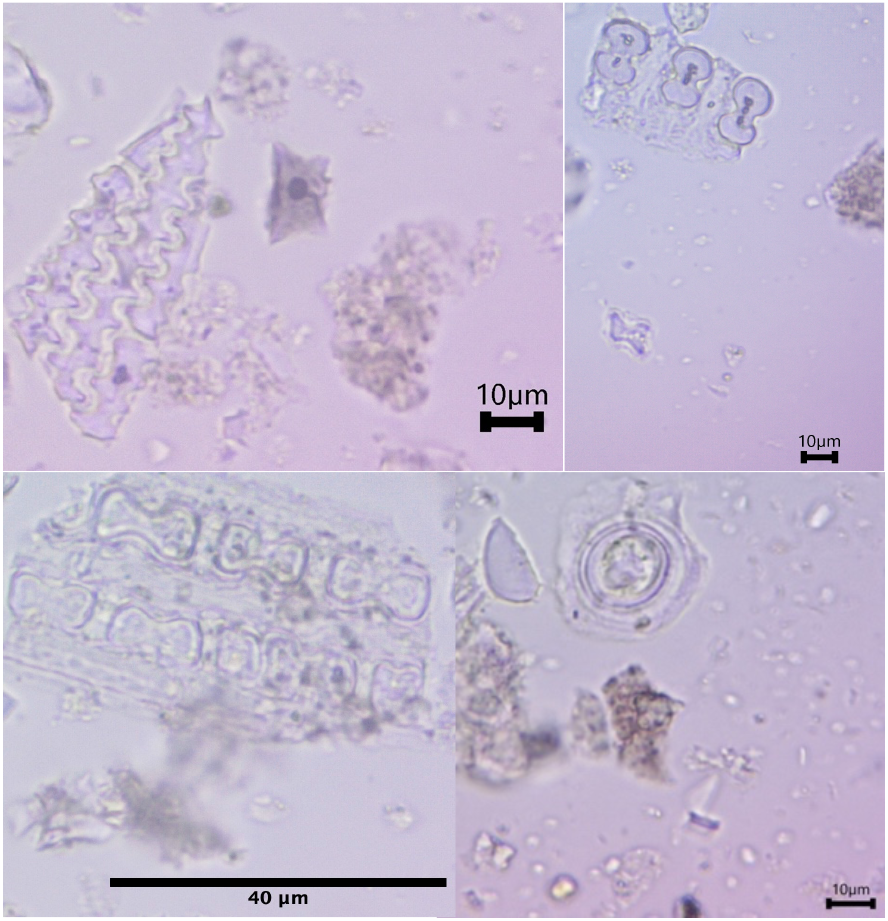
Phytoliths from layers 10 and 11 (from top left to bottom right : Elongate sinuate (ELO_SIN) and saddle (SAD), Bilobate (BIL) and cross (CROSS), skeletons with bilobates, Trichome (Hair base).

Intra-assemblage variability is evident within these units. Elongate dendritic (ELO_DEN) morphotypes are particularly abundant in Sample 6 (SU 11a) and Sample 7 (SU 10). Saddle (SAD) morphotypes occur in Samples 4 and 5 (SU 11b–11a transition), but are absent in Samples 6 and 7.

Intra-assemblage variability is evident within these units. Elongate dendritic (ELO_DEN) morphotypes are particularly abundant in Sample 6 (SU 11a) and Sample 7 (SU 10). Saddle (SAD) morphotypes occur in Samples 4 and 5 (SU 11b–11a transition), but are absent in Samples 6 and 7.

A substantial proportion of indeterminate phytoliths was also recorded, including fragments consistent with trichomes, stomata, and tracheary elements. Several specimens preserve partial anatomical connections (“phytolith skeletons”), especially in Samples 6 and 7.

### Microbotanical residues and use-wear analysis

Nine lithic artefacts were examined. Only one specimen (N160) yielded well-preserved residues suitable for detailed analysis.

#### Tool N160 (Layer 10, 8258–4007 cal BP)

N160 is a cobble fractured/splitted by anvil percussion and subsequently retouched unifacially on its non-cortical (convex) surface (Fig. 5). Use-wear analysis indicates both longitudinal and transverse motions along the active edge. White residues are concentrated along the working edge, particularly between retouch scars. Microscopic observation revealed preserved cell-wall structures consistent with monocot tissues, comparable to bamboo (Fig. 6).

**Figure 5.**
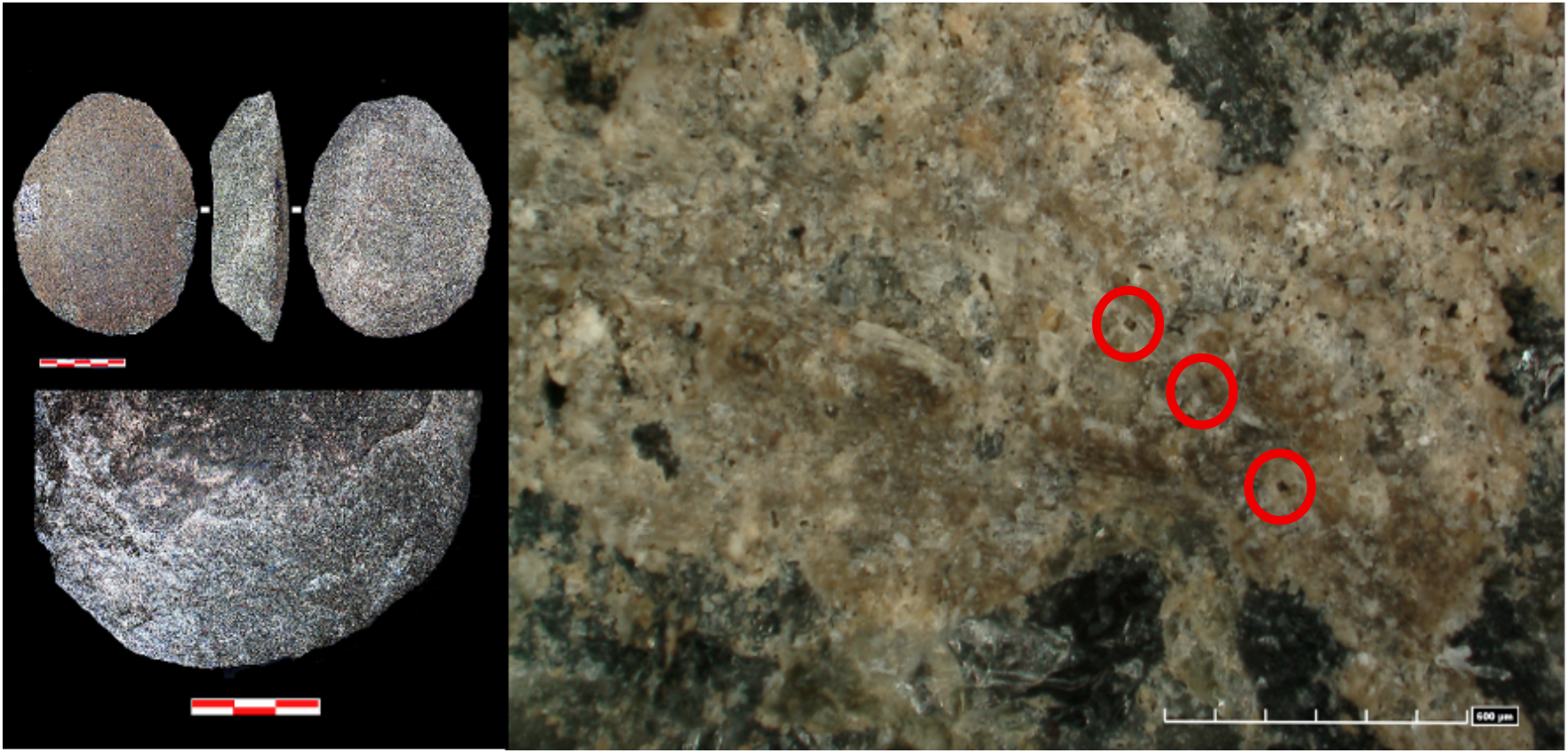
Unifacial cobble tool no. 160 (after I. Clemente-Conte), with a microphotograph of a monocot epidermis showing parallel bands and short cells (red circles) preserved in anatomical order on its surface.

**Figure 6.**
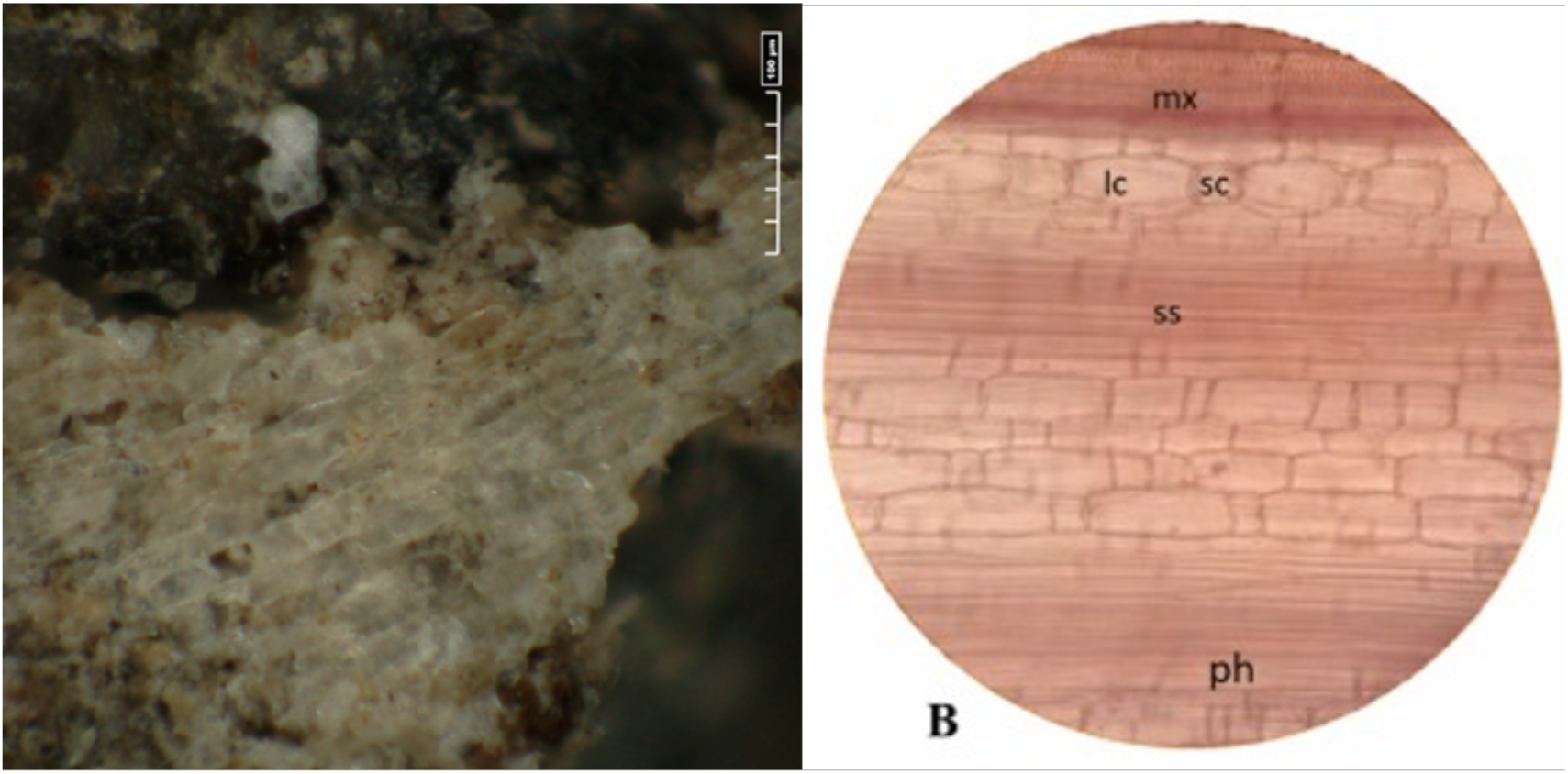
On the left, micro-plant remains of a monocot leaf–culm are preserved on the active surface of lithic tool N160, showing cells with a parallel-band arrangement. On the right, tangential longitudinal section of the pith of *Bambusa vulgaris* var. *striata* (magnification 10 × 40) from Mulyaningsih et al. (2022): lc, long cell; mx, metaxylem; ph, phloem; pp, pith parenchyma; sc, short cell; ss, sclerenchyma sheath.

#### Tool LS13 A29 n°27 Z1= 83 (Hoabinhian Stratigraphic Unit, Layer 9, room2, Unit I) - from I. Clemente-Conte’s report

Three additional artefacts (LS13 N27, LS17 B35 N24, LS16 C34 N58) exhibited degraded micro-plant residues with only partially preserved anatomical features, which, when considered alongside the functional interpretation of the tools, may tentatively be compatible with bordered pits indicative of coniferous wood. However, these remains are too poorly preserved to support a secure taxonomic identification (Fig. 7).

**Figure 7.**
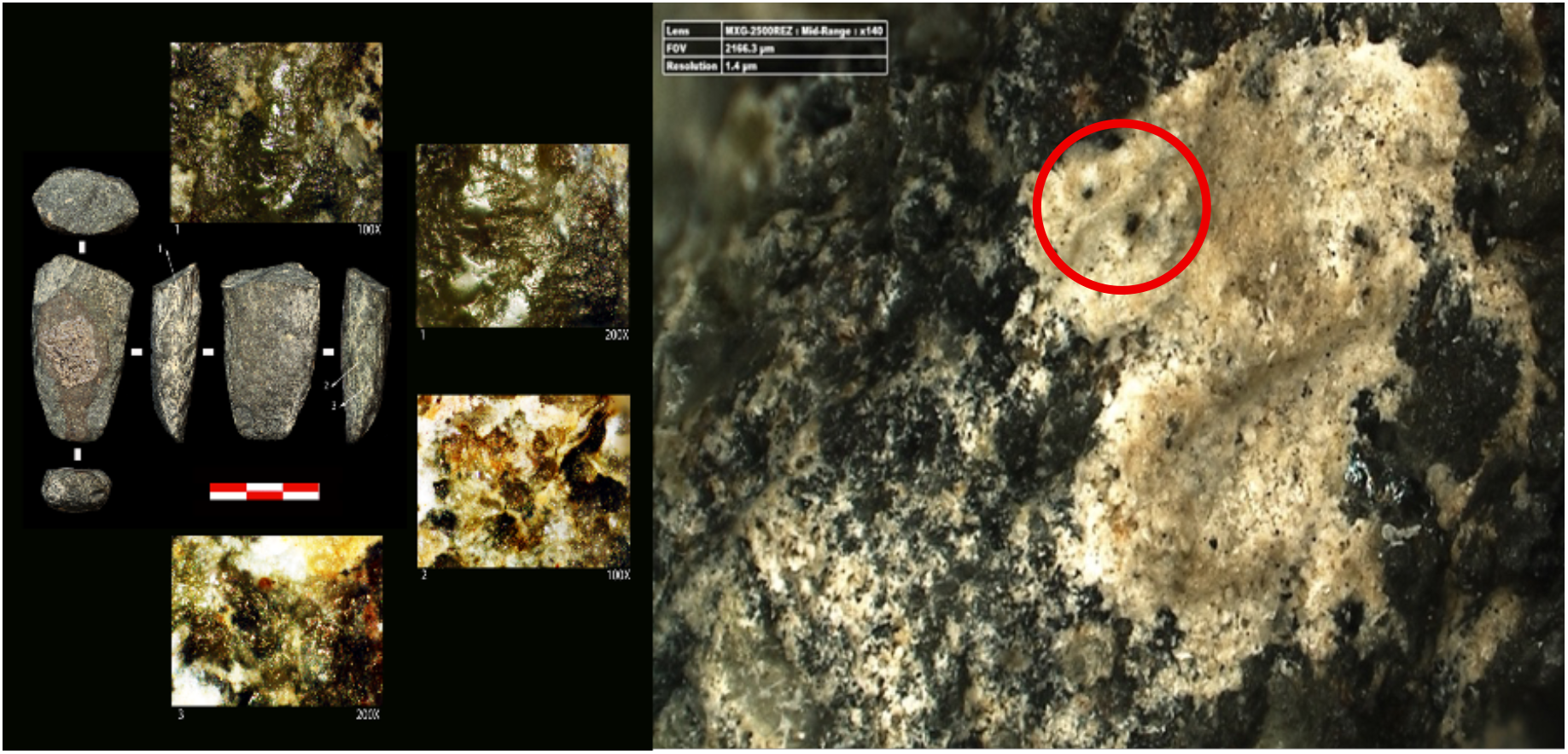
Plant pits, perhaps bordered pits (red circle) and tentatively suggestive of conifer tissue, are present, although preservation is poor. This tentative identification is compatible with the use-wear observations and the overall tool morphology, which together may indicate a hafted lithic implement.

#### Other artefacts

Three artefacts (LS13 N27, LS17 B35 N24, LS16 C34 N58) yielded poorly preserved residues. These show limited anatomical features that may be compatible with bordered pits, but preservation is insufficient for secure identification (see appendices).

## Discussion

The phytolith assemblage from Laang Spean suggests a diachronic shift between SU 9 and SU 10–11, although the available data do not allow a detailed reconstruction of local environmental change. In SU 9, the phytolith assemblage is strongly dominated by palm taxa, a pattern consistent with other sites from the same period and region. At Xincun in southern China, for example, Yang et al. (2013) and Wang (2024) identified abundant palm phytoliths, probably including sago palm, together with banana, water chestnut and fern remains, suggesting exploitation of diverse wet and forested habitats. This pattern aligns with present-day deciduous dipterocarp forest, where palm patches occur locally (Maxwell, 2004).

The grass-rich assemblages in SU 10–11 are notable for their diversity and density. Elongate dendritic morphotypes and, in some samples, saddle forms were recorded, the latter sometimes associated with Chloridoideae grasses. Their occurrence could be consistent with the presence of grasses adapted to warm and seasonally dry environments, but this interpretation remains tentative, as phytolith distributions are shaped by both ecological and taphonomic factors. The predominance of vegetative phytoliths over seed-related morphotypes may also partly reflect the known tendency of grasses to produce and preserve leaf and culm forms more readily than reproductive tissues (Albert & Weiner, 2001; Cabanes, 2020).

Residue analysis on lithic artefacts also supports the hypothesis of a largely “lost” plant technology, in which organic implements are rarely preserved as artefacts or raw materials but remain indirectly visible through the residues they leave on stone manufacturing tools. Such missing implements were probably used as hunting weapons or as components of composite tools with organic hafts, and would have been lighter and more portable than cobble tools, while no less functionally effective for specific tasks (Forestier, 2003; Sellato, 2021; Forestier et al., 2026). Here the residue analysis of lithic artefacts provides complementary, though limited, evidence as only one artefact, N160 (from layer 7), yielded residues sufficiently preserved for detailed observation. It contains microremains consistent with monocot leaf–culm tissue, most likely derived from tissues close to the pith, as suggested by anatomical characteristics that closely match bamboo tissues in this zone and point to intensive working and shaping (Fig. 5 & 6). The sediment can therefore be read, at least in part, as providing a strong signal of plant processing for technological purposes in a cave context, where activities such as hafting, shaft manufacture, and mat or cordage production would generate abundant vegetative debris. The identified scraping activity is consistent with this interpretation, although a dietary role cannot be excluded: the tool may also have been used to remove the plant epidermis for consumption, and both functions could have been performed using the same implement. Further ethnoarchaeological work with a range of monocot taxa is needed to refine these interpretative possibilities.

Beyond monocot seed-like exploitation, the management of arboreal taxa within agroforestry-like systems appears at least as complex and clearly requires interdisciplinary approaches that can cross-reference different categories of plant remains. Maxwell’s work (1999; 2004) tracking anthropogenic fire events demonstrates the difficulty of identifying clear human impact markers in vegetation records. Nevertheless, these fire events can be interpreted as forest expansion (Kealhofer, 2003) and soil management strategies in monsoonal climate for agroforestry management purposes (Maxwell, 2004; White et al. 2004). Mourer’s (1977) pollen and spore study already revealed a diverse arboreal community at Laang Spean, including Anacardiaceae, Sapotaceae, *Bombax* sp., *Ceiba* sp. and *Duabanga* sp., alongside Poaceae, a pattern that aligns well with the phytolith results presented here. Taken together, these data point to the use of structurally complex forest–fluvial mosaics, where nuts, fruits, tubers and grass seeds formed an integrated resource base.

Several limitations should therefore be noted. However, there are clear limits to what phytolith alone can reveal about arboreal taxa, especially in the absence of a local reference collection to refine the identification of epidermal phytoliths (hairs, stomata, etc.). The trichomes, stomata and phytolith “skeletons” observed in the assemblage strongly support both the functional interpretations and the presence of multiple plant taxa, but taxonomic identifications must be grounded in regionally appropriate comparative material. For example, stomata (microscopic pores on leaf surfaces that regulate gas exchange) were identified in this assemblage but could not be assigned to specific plant groups due to the absence of such a reference framework. As stomatal morphology, including shape and spatial arrangement, varies across taxa, it holds significant taxonomic potential.

In future work, we aim to systematically document plants associated with particular activities and to combine this with the analysis of the residues they leave on tools. This would allow for more precise identification of taxa, the plant parts processed, and the intensity of use. Grounded in comparative anatomy and histology, this approach would significantly improve the interpretative resolution of microbotanical data. It would also help distinguish different functional signals within mixed assemblages, including technological versus dietary uses.

## Conclusion

This study demonstrates that combining phytolith assemblages with micro-plant residues on lithic tools refines interpretations of Hoabinhian plant use and “forest technologies” at Laang Spean. Stratigraphic phytolith data show a clear contrast between hoabinhian phases, with the stratigraphic unit (layer 9) dominated by palm phytoliths and layers 10 and 11 characterised by higher phytolith densities and diagnostic grass forms, alongside diverse arboreal morphotypes.

Although micro-plant residues were rare on the studied lithic tools, one artifact preserved leaf-culm tissues attributable to monocots, most probably bamboo, directly linking specific tool use to the processing of flexible plant stems. Together, these lines of evidence suggest that Hoabinhian communities at Laang Spean exploited a combination of palm tree, grass and arboreal resources within a complex forested environment, and highlight the value of integrated anatomical and phytolith approaches for reconstructing the material and environmental dimensions of Pleistocene–Holocene forager lifeways in Southeast Asia as a civilisation of « Plant-life » (Gourou, 1948; Solheim, 1972).

## Appendices

*Tool LS_17_Ab35_n°24_5*.*3 Z1=90 cm (Hoabinhian Stratigraphic Unit, Layer 9, room2, Unit I) from I*.*Clemente-Conte’s report*

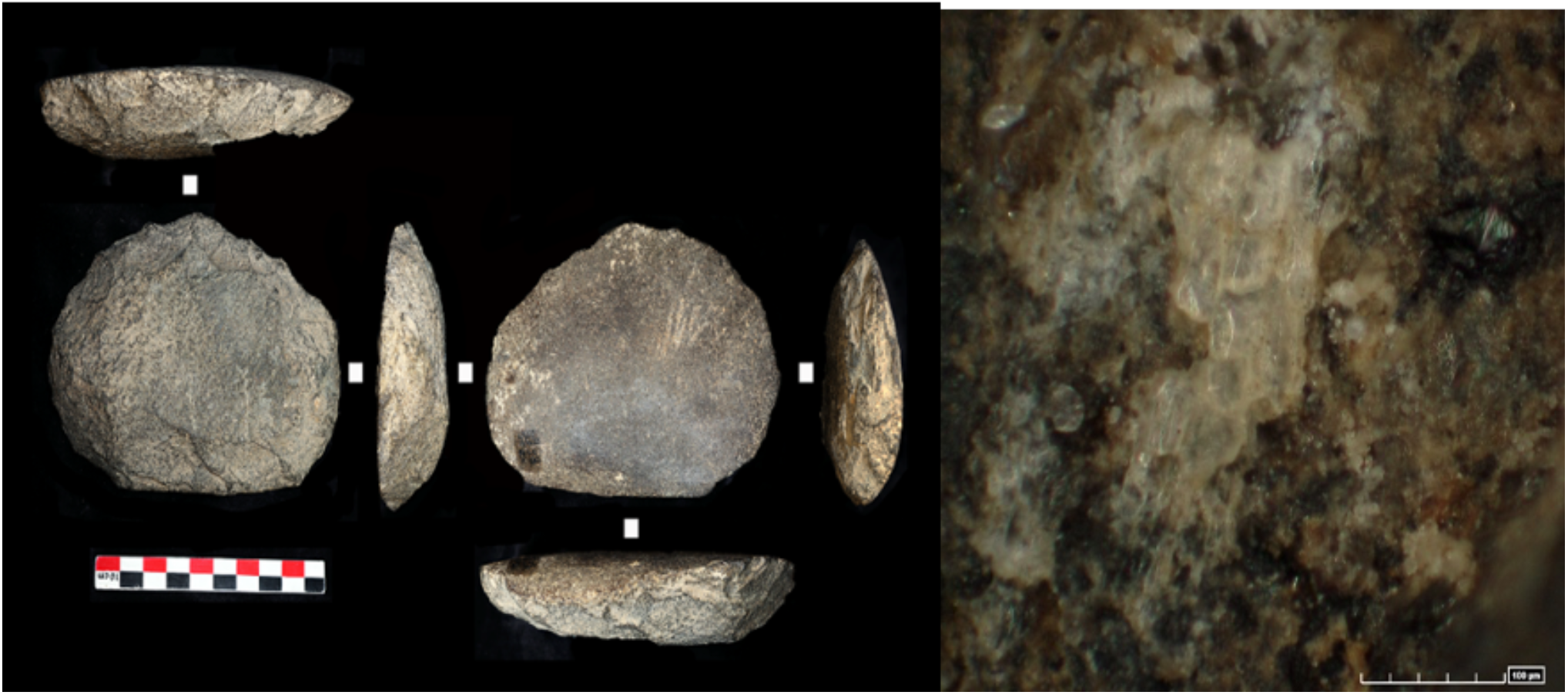

Plants tissues are present, but preservation is poor.

*Tool LS16 C34 n°58 Z1=66 ((Hoabinhian Stratigraphic Unit, Layer 9, room2, Unit I) from I*.*Clemente-Conte’s report*

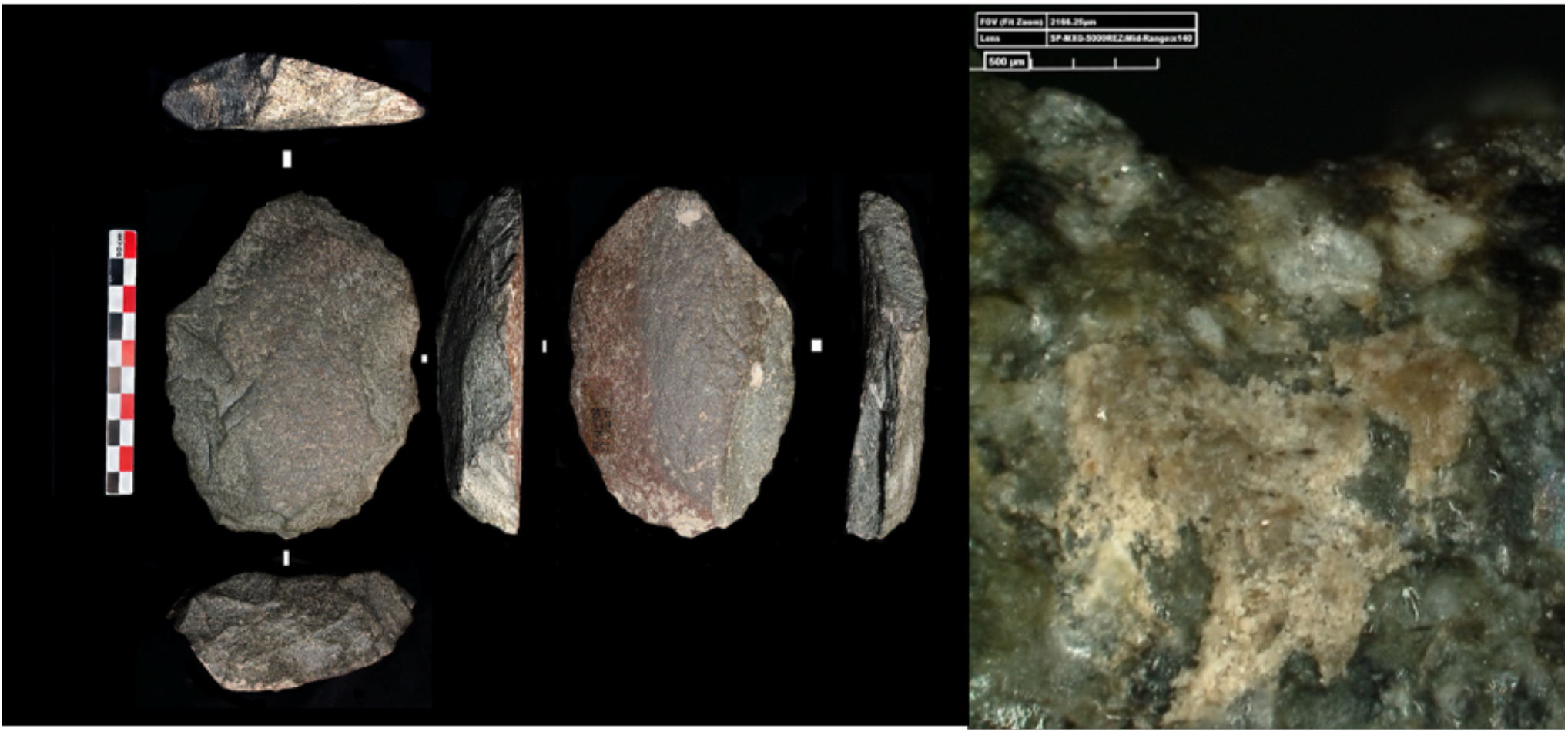

Plant tissues are present, but preservation is poor.

## Acknowledgements

Excavations at Laang Spean Cave were carried out by the “French Cambodian Prehistoric Mission” under the authority of the Ministry of Culture and Fine Arts of the Kingdom of Cambodia, in collaboration with the Royal University of Fine Arts (Phnom Penh). This research was supported by the Commission consultative des recherches archéologiques à l’étranger (Ministère de l’Europe et des Affaires étrangères, France), the Service de coopération et d’action culturelle (SCAC) of the French Embassy/Institut français du Cambodge (Phnom Penh), the Muséum national d’Histoire naturelle (MNHN), and UMR HNHP (CNRS).

We are grateful to the reviewers for their careful evaluation and constructive comments, which would prompt us to further refine and deepen the interpretation of our primary data.

## Data and supplementary information availability

Data are available online : https://doi.org/10.17605/OSF.IO/QP2W9 (Kerfant, C. E. (2026, July 25). The micro-plant remains preserved on lithics and in the associated sediment of the Hoabinhian Laang Spean cave site, Battambang province, Cambodia.);

Supplementary information is available online: https://doi.org/10.17605/OSF.IO/DVZ9E (Kerfant, C. E. (2026, July 25). Supplementary Materials. Retrieved from osf.io/dvz9e).

